# The Cognitive Critical Brain: Modulation of Criticality in Task-Engaged Regions

**DOI:** 10.1101/2023.06.29.547080

**Authors:** Xingyu Liu, Xiaotian Fei, Jia Liu

**Author notes:** Co-first authors (Xingyu Liu, Xiaotian Fei).

## Abstract

The constantly evolving world necessitates a brain that can adapt and respond to rapid changes. The brain, conceptualized as a system performing cognitive functions through collective neural activity, has been shown to maintain a resting state characterized by near-critical neural activity, poised to respond to external stimuli. The dynamic adaptation of near-criticality during various tasks, however, remains poorly understood. In this study, we utilized the prototypical Hamiltonian Ising model to investigate the modulation of near-criticality in neural activity at the cortical subsystem level during cognitive tasks. Specifically, we theoretically simulated cortical 2D-Ising models *in silico* using structural MRI data and empirically estimated the system state *in vivo* using functional MRI data. First, our findings corroborated previous studies that the resting state is typically near-critical as captured by the Ising model. Notably, we found that cortical subsystems changed their criticality levels heterogeneously during a naturalistic movie-watching task, where visual and auditory cortical regions were fine-tuned closer to criticality. A more fine-grained analysis of the ventral temporal cortex during an object recognition task revealed that only regions selectively responsive to a specific object category were tuned closer to criticality when processing that object category. In conclusion, our study supports the *cognitive critical brain hypothesis* that modulating the criticality of subsystems within the hierarchical modular brain may be a general mechanism for achieving diverse cognitive functions.

## Introduction

The fundamental goal of neuroscience is to understand how the brain achieves complex behaviors with robustness and flexibility. One approach is to conceptualize the brain as a system realizing diverse cognitive behaviors through collective phenomena emerging from myriads of interconnected neurons. The *critical brain hypothesis* [1], which posits that a healthy brain operates in a state of criticality, has gained prominence in recent years. Criticality, an overarching concept in statistical mechanics and complex science, refers to a unique system state perching between distinct phases or regimes, such as ordered and disordered phases (Fig. 1A), where the system’s symmetry is broken. This state facilitates the emergence of long-range correlations and fluctuations, leading to distinct macroscopic behaviors absent in individual phases or regimes. Consequently, criticality is considered the point where macroscopic complexity arises from microscopic simplicity [2][3]. To date, criticality has proven effective in understanding various systems, including natural and social systems [4][5], and the brain is no exception. Empirical evidence supporting the *critical brain hypothesis* has shown that the resting brain, characterized by unconstrained spontaneous activity, resides near-criticality [6][7][8][9][10]. However, the cognitive significance of the critical brain remains unresolved, as there is no clear connection between criticality and versatile cognitive functions. In this study, we asked whether criticality serves as a governing principle for the cognitive brain, enabling the relatively structure-fixed brain to functionally reconfigure itself to perform behaviorally relevant computations and accomplish various tasks immanent in a natural environment.

**Fig. 1.**
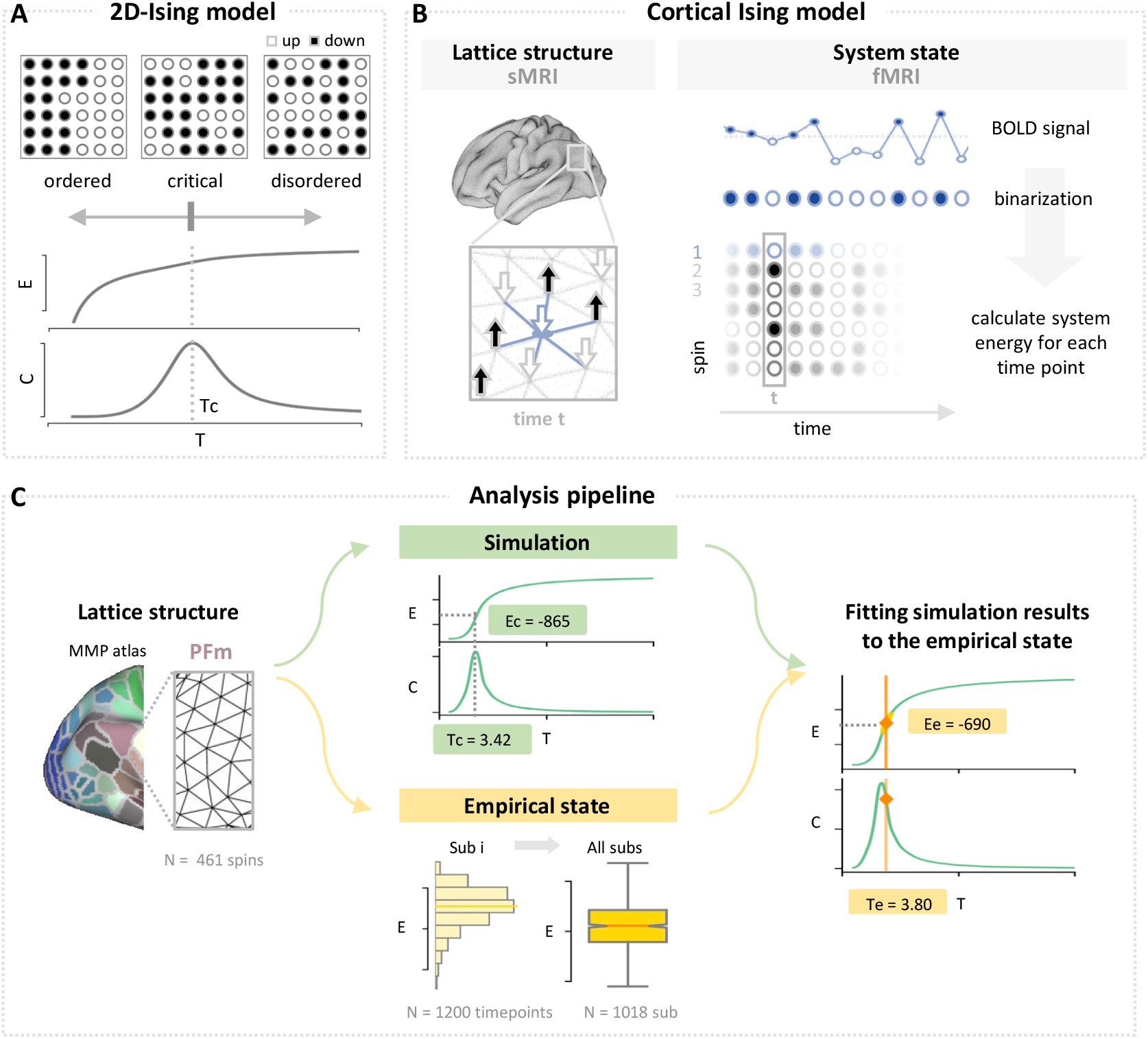
Schematic illustration of modeling the cortical Ising system. **(A)** 2D-Ising model. Example lattice structure and configurations (top), and thermodynamic properties including specific heat (middle) and energy (bottom) of the 2D-Ising model at three featured temperatures: *T < Tc*, ordered state; *T* = *Tc*, critical state; *T > Tc*, disordered state. **(B)** Modeling cortical Ising system using sMRI and fMRI data. Left panel: the lattice architecture information required for cortical lattice structure is provided by sMRI. Right panel: the cortical dynamics provided by fMRI data can be converted to spin activity through binarization. **(C)** Analysis pipeline taking PFm subsystem as an example. Left: the PFm parcel and lattice architecture information from the MMP atlas. Middle top: simulations at various temperatures generated an energy curve and a specific heat curve. The critical state was determined by the peak of the specific heat. Green textboxes: the critical temperature and energy (*Ec*). Meddle bottom: Estimation of empirical temperature (*Te*) and energy (*Ee*) of cortical Ising system using fMRI data. The empirical energy was determined by the median energy across all timepoints for each individual and then across all participants. Right: Fitting the simulation results to the empirical data revealed a near-critical state of the cortical Ising system-PFm. The energy and specific heat as a function of temperature from the simulation are plotted in green lines; the empirical states are marked in orange and annotated in textboxes.

An intuitive method to examine neural dynamics under tasks is to analyze neurophysiological recordings from neurons. This method has found that neural activity maintains or approaches criticality during cognitive tasks [11][12], for instance, in turtle’s visual cortex after perturbation induced by external stimuli [13] and in mouse’s cortical layers 2/3 during conditional learning [14]. However, this method is limited by a sparse sample of neurons from a small region of the brain, potentially leading to a subsampling issue and therefore inconsistent results [15][16]. Indeed, several studies using this method have reported deviations from criticality [17][18]. To obtain a more global perspective, researchers have utilized whole-brain imaging techniques, such as functional magnetic resonance imaging (fMRI), electroencephalogram, and magnetoencephalography. Nevertheless, the findings remain contentious, with some reporting self-organized criticality during a visual detection task [19] and near-critical state of neural avalanches during visual recognition [20], while others observing shifts away from criticality during a task requiring higher attentional load [21]. This inconsistency may arise from treating the brain as a homogeneous system, neglecting the diverse involvement of distributed cortical regions (i.e. subsystems) in different tasks [22][23][24]. Therefore, it is crucial to prioritize spatial resolution to accurately capture cortical heterogeneity. The unparalleled solution lies in utilizing structural MRI (sMRI) and fMRI to discern structural and functional heterogeneity, respectively, thus accounting for the homogeneity within local subsystems and the heterogeneity among them.

Specifically, we utilized the criticality of Hamiltonian systems within the framework of statistical mechanics to directly assess whether the brain undergoes a phase transition. The prototypical Hamiltonian lattice system is the Ising model, which assumes nearest-neighbor interactions. The cortical surface, the outer layer of the brain mainly responsible for cognition, is a sheet-like structure with distinct geometry, rendering it suitable for a spin lattice of a 2D-Ising model (Fig. 1B). Therefore, using detailed cortical anatomy provided by sMRI and dynamic neural activity provided by fMRI, we modeled cortical subsystems as 2D-Ising spin lattice systems with high biological fidelity, with their states directly characterized by thermodynamic quantities. In line with previous studies, we found that the resting brain exhibited near-criticality as characterized by the Ising model, which persisted across cortical subsystems as well. Under the tasks of a naturalistic movie watching and well-controlled object recognition, we observed a heterogeneous functional modulation on criticality for cortical subsystems, demonstrating a strong association between brain state fine-tuning towards criticality and functional engagement. Collectively, these findings suggest that subtle modulation of criticality serve as a general mechanism supporting the cognitive division of labor in realizing a diverse function repertoire.

## Results

### Cortical subsystems operate at near-criticality during rest

We first characterized the empirical cortical state of the resting brain to establish a baseline for task modulation. This was accomplished by directly measuring the control parameter of the cortical Ising model, temperature, during rest. To this end, we reconstructed the lattice architecture derived from the group-averaged surface geometry of the bilateral hemispheres using sMRI data from the Human Connectome Project (HCP) [25], resulting in dense triangular lattice architectures comprising over 30,000 vertices (i.e. sites) per hemisphere. We then obtained the brain subsystems, where Ising model simulations were performed, using the canonical multi-modal parcellation (MMP), which partitions the bilateral hemispheres into 180 regions based on multiple anatomical and functional features [26]. For instance, the lattice architecture of subsystem PFm in the parietal lobe (Fig. 1C left panel) was derived from the surface geometry of the PFm parcel from the MMP, which contains 461 vertices with known neighborhood relationships. We acquired the energy and specific heat curves for the Ising system-PFm through simulation, with temperature ranging from 0.001 to 1000, using a Wang–Landau sampling implementation [27][28]. The critical point, where specific heat diverges, was identified at the peak of specific heat curve as the center of the criticality zone, with the corresponding temperature and energy being the critical temperature (*Tc*) and critical energy (*Ec*). Temperatures falling within the half width at half maximum of the specific heat was considered near-criticality. After obtaining the theoretical criticality indexed by *Tc* and *Ec*, we assessed the state of the human brain during rest using functional neural activity from resting state fMRI (rs-fMRI). As shown in Fig. 1B, the neural activity was binarized for each vertex along the temporal domain, with above-average activity representing spin-up and below spin-down. The empirical energy (*Ee*) defined by the Ising-format Hamiltonian was quantified at each time point of the rs-fMRI, which was used to derive the empirical temperature (*Te*) by fitting the simulated energy to the *Ee*. Consequently, the *Te* of the Ising model-PFm was found to be 3.80, within the range of near-criticality centering on the *Tc* (3.42) (Fig. 1C right panel), suggesting that the PFm was in a near-criticality state during rest.

We extended our analysis to encompass all 180 brain regions of the entire cerebral cortex. As illustrated in Fig. 2A, critical or near-critical states were observed across all subsystems, consistent with previous studies [6][7][8]. To quantitatively evaluate the difference between Te and Tc, we calculated the temperature difference ratio, defined as (*Te − Tc*)*/Tc*, for each subsystem. Relatively small ratios (Mean = 11.75%) were found for all subsystems in both left and right hemispheres (Fig. 2B), with all Te values marginally higher than *Tc*, suggesting a slight intrusion towards the disordered phase. Interestingly, despite the prevalent near-criticality across all subsystems, minor variations among subsystems were observed (SD = 3.72%), suggesting heterogeneity in near-criticality across subsystems. Specifically, cortical subsystems located in the intraparietal, supplementary motor area, and posterior medial cortex corresponding to the multiple demand network [29][30], exhibited smaller temperature difference ratios, implying a closer proximity to criticality (Fig. 2B).

**Fig. 2.**
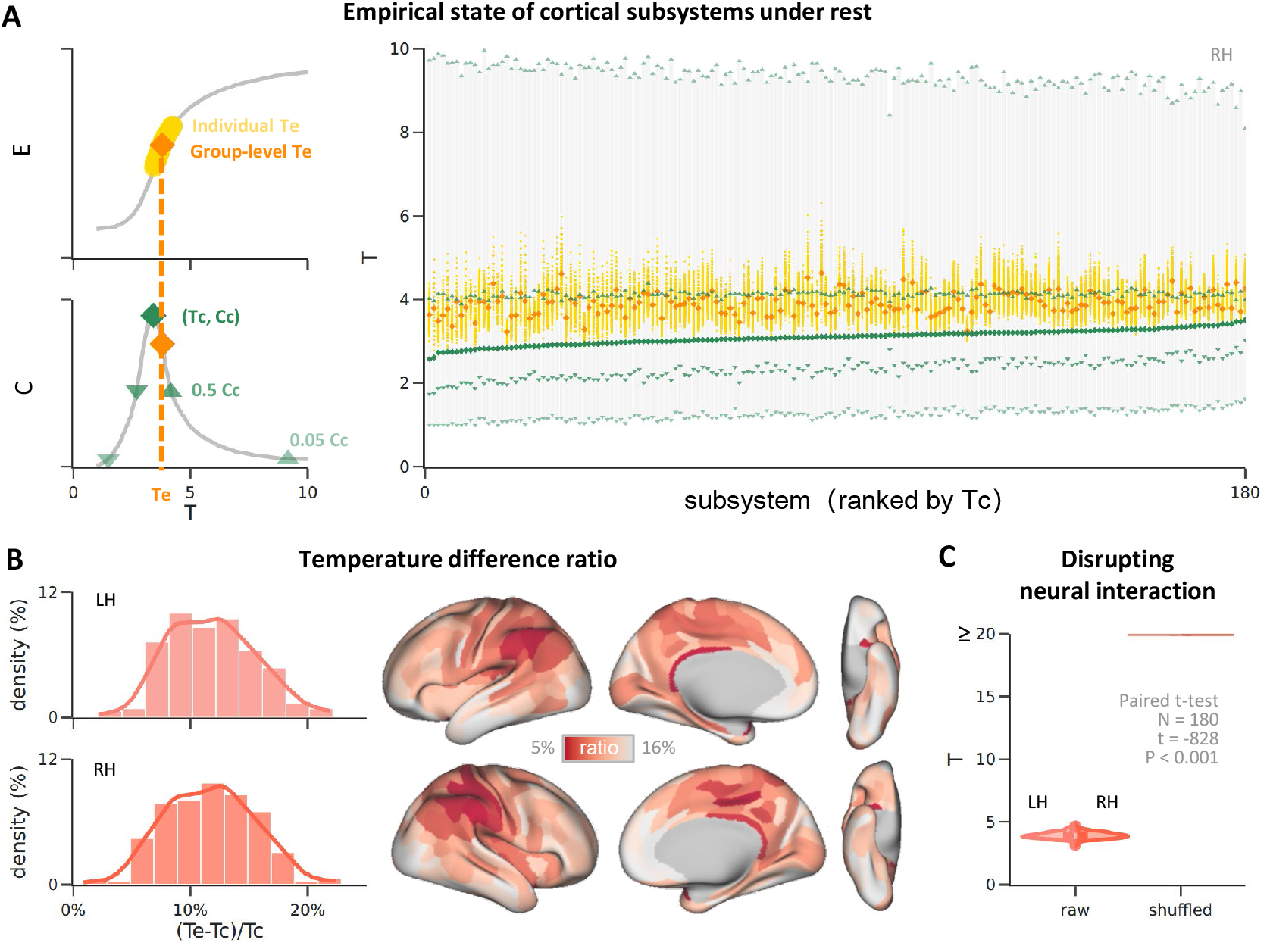
All cortical subsystems operated at near-criticality during rest. **(A)** Estimated empirical temperature (*Te*) and simulated critical temperature (*Tc*) for all 180 brain region subsystems from the right hemisphere. The x-axis is sorted by *Tc*, and the legend for the right panel is provided in the left panel. *Cc*: the specific heat of the critical point. **(B)** A small temperature difference ratio was revealed in all brain regions subsystems in both hemispheres. Left: Distribution of temperature difference ratio, defined as (*Te − Tc*)*/T c*, across all brain regions. Right: brain map of the temperature difference ratio. **(C)** Estimated Te of the original fMRI data and shuffled fMRI data. The null model using temporally shuffled fMRI data that damages the neural interaction showed a disordered state far from Ising criticality.

To rule out the possibility that the near-criticality during rest was merely a byproduct of lattice architecture or certain characteristics inherent to fMRI modality, we disrupted the Ising-format interactions between spins while preserving their individual statistical properties by temporally shuffling each spin and then measuring the temperature of this shuffled fMRI data. This control analysis revealed a disordered phase significantly distanced from criticality in all subsystems (Fig. 2C), confirming that the resting cortical subsystems were indeed at near-criticality. Besides, the minimal temperature fluctuation over various time periods adhered to the equilibrium assumption of the Hamiltonian system (Supplementary Figure 1), a prerequisite for the application of Ising models in criticality measurement.

Notably, the near-criticality of the resting brain was not only prominent at the level of cortical subsystems, but was also evident at larger scales. Indeed, similar results of near-criticality during rest were found at the brain network level (e.g. visual, attention, and default mode networks) and the whole hemisphere level (either left or right hemisphere) using the conventional Metropolis Monte Carlo algorithm (Supplementary Figure 2). Taken together, the subsystems of the resting brain operated at a state of near-criticality, albeit with a slight variance in their inclination towards the disordered phase.

### State criticality shift correlates with functional engagement across cortical subsystems

Having established the near-criticality of the resting brain as a baseline, we investigated how criticality is modulated to accommodate varying cognitive demands. Given the functional segregation of the human brain and heterogeneity during rest, we expected that task-induced changes in criticality would vary across subsystems. To this end, we utilized task-fMRI (t-fMRI) data from participants engaged in a movie-watching task, which is known to elicit a broad range of cognitive functions and recruit disparate cortical regions throughout the brain [31][32].

Utilizing a separate dataset collected in a 7T MRI scanner [25], we first replicated our previous finding with different spatial and temporal resolutions that the resting brain operated at near-criticality (Supplementary Figure 3). Importantly, we found that the task-engaged brain during moviewatching also exhibited near-criticality across all subsystems, with a similar tendency towards the disordered phase Fig. 3A. The findings that pertain to both the resting and task-engaged brain suggest that the cortical system maintains a stable state of near-criticality, and that any potential task-induced modulations of criticality likely remain subtle within the critical zone, rather than involving drastic transitions from one phase to another.

**Fig. 3.**
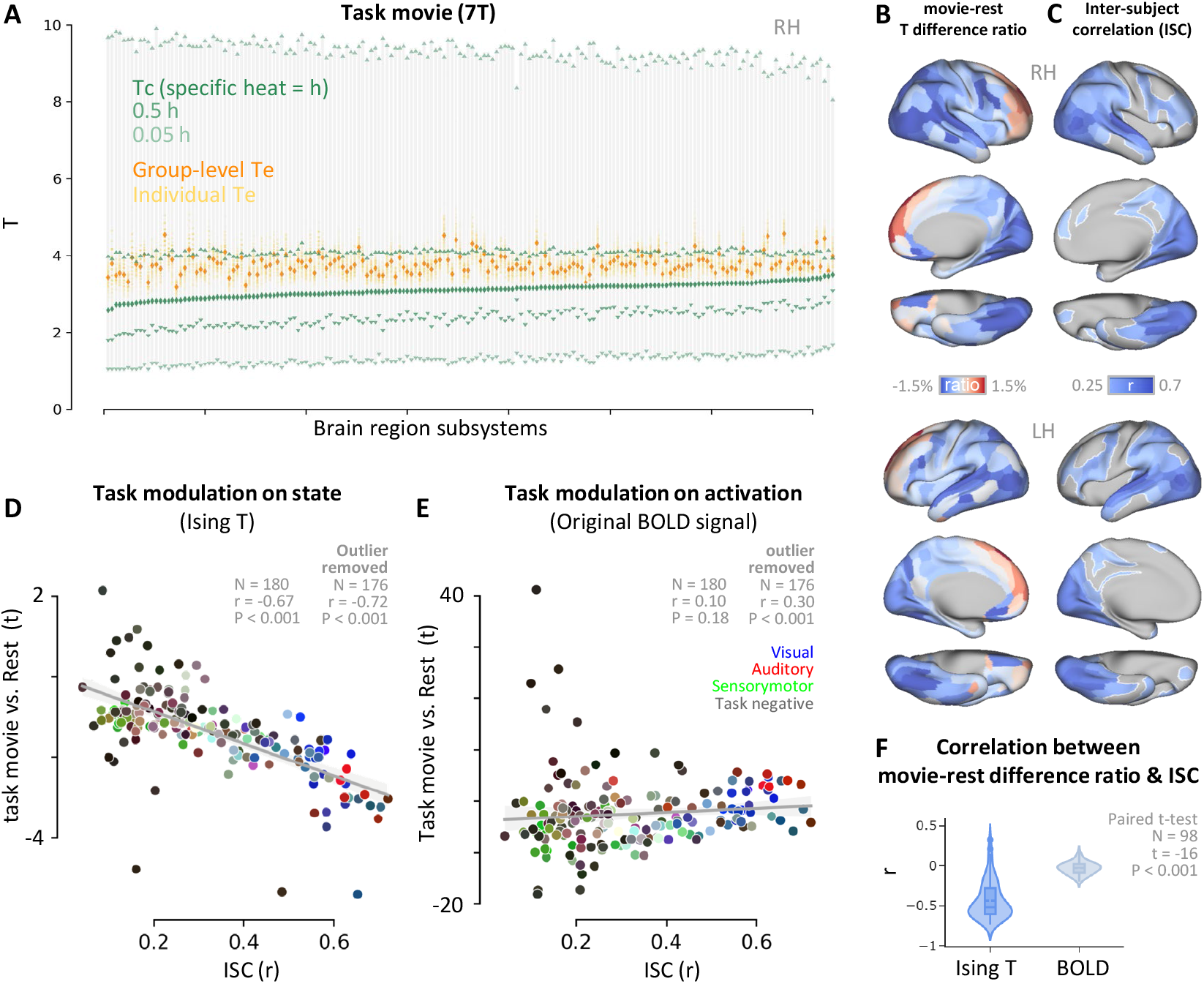
Functional engagement shifts cortical subsystems toward criticality under a naturalistic movie-watching task. **(A)** The same plot as in Fig. 3A for the movie-watching task using data collected in 7T MRI scanner. Near-criticality was found for all cortical region subsystems under the task. **(B)** The t-value map of task movie versus rest. A paired t-test was performed between the empirical temperature estimated from the movie-watching state and resting state for each cortical region subsystem. **(C)** Inter-subject correlation map of the movie-watching task. **(D)** Scatter plot of the task modulation on criticality attunement (task versus rest of Ising energy) and functional engagement (ISC). Each point denotes a subsystem. **(E)** The same plot as in (D), but with the y-axis replaced by the task modulation on raw activation. The values of the corresponding subsystems in both hemispheres were averaged in (C) and (D). **(F)** Bar plot of individual correlation values between difference ratios of Ising temperature/original BOLD signal and ISC.

To examine functional modulations on criticality at the subsystem level, we compared the criticality of each subsystem during the task to that during rest, with an index of a temperature difference ratio, defined as (*Te*_*movie*_ – *Te*_*rest*_) / *Te*_*rest*_. As illustrated in Fig. 3B, the effect of functional modulations on criticality varied across subsystems. Some subsystems located in the occipital, temporal, and parietal lobes exhibited lower temperatures during the task than at rest. The decrease in temperature indicates finetuning towards criticality. Conversely, certain subsystems in the frontal lobe displayed a minor shift towards higher temperatures, slightly moving away from criticality. Detailed information on cortical subsystems that shifted closest to (temperature difference ratio < -2.4%) or furthest from (> 1%) criticality is listed in Supplementary Table 1.

To further investigate the functionality of task modulation on criticality, a visual inspection revealed that subsystems fine-tuning closer to criticality aligned with cortical regions engaged in the movie-watching task reported previously [33][34][35]. To quantify this observation, we measured the functional engagement of subsystems during movie-watching using a cognitive neural benchmark, intersubject correlation (ISC) [36], which captures the synchronization of neural signals across participants in response to identical stimuli [37] [38]. Consistent with previous studies[39], subsystems showing moderate to high ISC were primarily involved in processing visual, auditory, language, and event-related information (Fig. 3C), all essential functions for the movie-watching task.

Crucially, we evaluated the relationship between the effect of task modulation on criticality (indexed by the contrast of *Te*_*movie*_ versus *Te*_*rest*_) and functional engagement (indexed by ISC). We found a strong correlation between criticality fine-tuning and ISC (r = -0.69, P < 0.001) (Fig. 3D), suggesting that subsystems that fine-tuned more towards criticality during the task were more engaged in the task. Conversely, when the collective activity of interacted neurons, which underpins criticality, was averaged as a univariate response magnitude, the effect of task modulation (indexed by the contrast of response magnitude during the task versus that during rest) showed nil or, at most, weak correlation with the ISC (r = 0.09, P = 0.22; r = 0.30, P < 0.001 when outliers > 3 IQR excluded) (Fig. 3E). Therefore, dynamic changes in criticality during tasks are likely a more sensitive neural signature for task engagement than the changes in univariate activity magnitude traditionally relied upon in analyses. In addition, this pattern was also observed at the individual participant level, where in each participant, the correlation coefficient between changes in criticality and ISC was larger than that between the univariate magnitude and ISC (Fig. 3F). Taken together, these findings suggest that dynamic changes in criticality during tasks may provide a means of realizing complex cognitive functions from seemly disordered activity of neuronal populations in the brain.

### Category-selective regions fine-tune towards criticality under structured visual task

The movie-watching task proves a naturalistic approach, revealing the effect of task modulation on criticality across diverse domains, spanning from primary sensory to high-level cognitive processing. Here, we further probed into a finer granularity within the domain of vision, where subsystems are tasked with processing highly specific tasks. To this end, we adopted an object recognition task consisting of four object categories (faces, bodies, tools, and places) (Fig. 4A), and examined the modulation on criticality of their corresponding object-selective cortical regions (i.e. ROI, regions of interest) (Fig. 4B) in the ventral temporal cortex (VTC). These four regions, derived from Cocuzza et al. [40] based on a large body of previous studies (Fig. 4B), include Face fusiform area (FFA) for faces, Extrastriate Body Area (EBA) for bodies, Lateral Occipital Complex (LOC) for tools, and Parahippocampal Place Area (PPA) for places. There were four levels of state criticality, each corresponding to the processing of one of the four object categories within their respective regions.

**Fig. 4.**
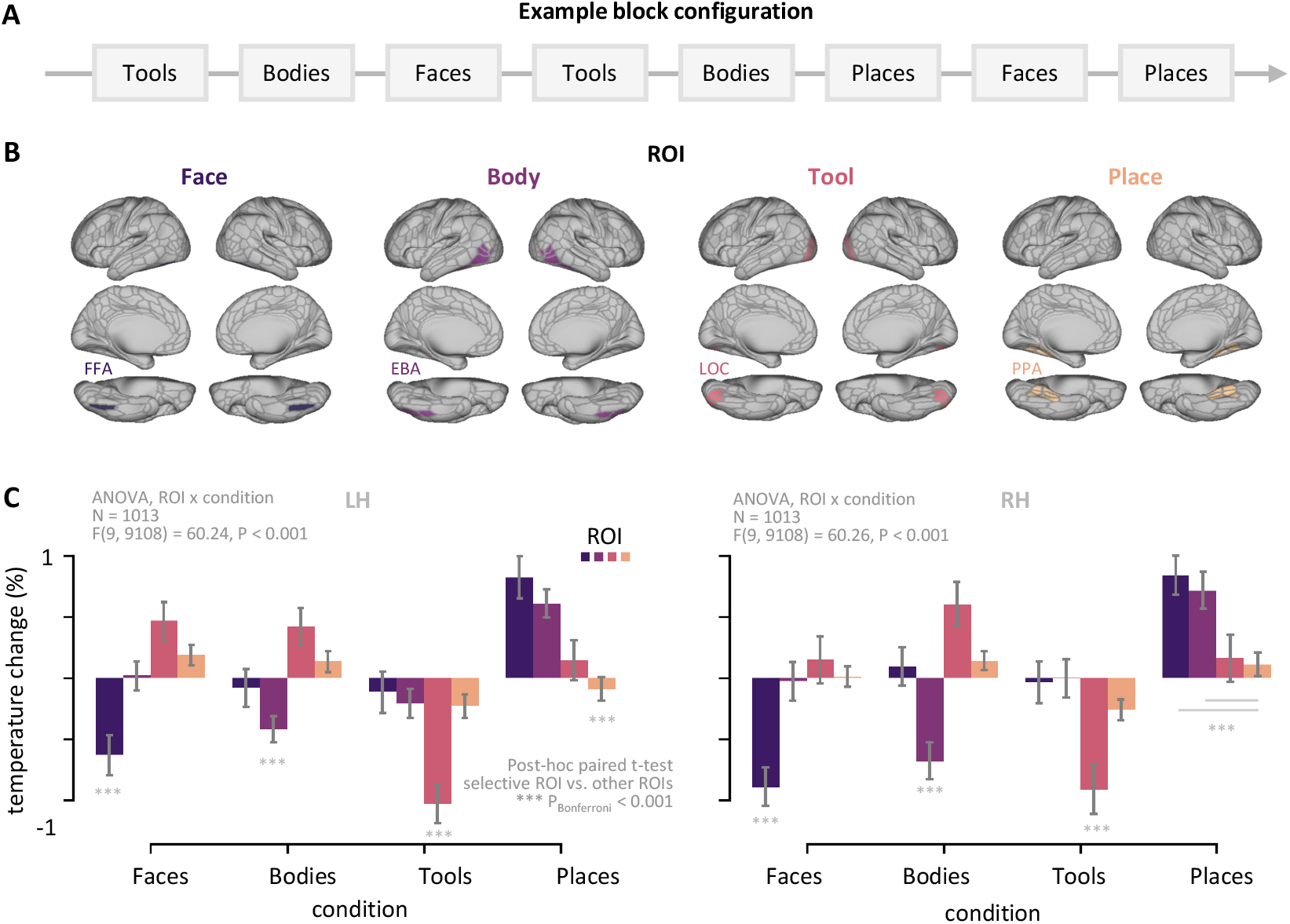
Object-selective subsystems in the VTC selectively fine-tuned towards criticality during object recognition. **(A)** Example block configuration of the task. Four object categories were presented including faces, bodies, tools and places. **(B)** ROIs, including the FFA, EBA, LOC and PPA, for the four categories. **(C)** Inter-subject correlation map of the movie-watching task. **(D)** Bar plot showing task modulation on system temperature across different ROIs under various category conditions. The values were aggregated across subsystems for each ROI. A significant interaction between ROI and category was observed in both hemispheres, supporting a locally domain-specific task modulation on brain criticality. LH: left hemisphere; RH: right hemisphere.

The task modulation on criticality was assessed by calculating temperature changes across all ROIs and all object categories. This analysis revealed heterogeneity in functional modulation on criticality in these regions for different object categories, with a significant interaction of ROI by object category in both hemispheres (F(9,9108) > 60, P < 0.001) (Fig. 4C), suggesting that the effect of task modulation on criticality in each region depended on the type of object category being processed. Post hoc statistical tests confirmed this observation, demonstrating a significant temperature decrease in an object-selective region (e.g. the FFA) when processing its preferred category (i.e. faces) (one-sample t-tests: all ts < 0, *P*_*Bonferroni*_ < 0.001). Moreover, this temperature decrease was most pronounced when compared to nonselective regions (i.e. EBA, LOC, and PPA) (paired t-tests: all ts < 0, *P*_*Bonferroni*_ < 0.001). That is, when a region conducted its designated task (e.g. the FFA in processing faces), this region fine-tuned towards criticality, characterized by a lowered temperature. Note that while among all ROIs tested, the lowest temperature in response to places was found in the PPA, the right PPA did not fine-tune toward criticality as compared to the resting state. This discrepancy could potentially be attributed to the functional heterogeneity of the PPA [41][42], supported by the inconsistent task modulation effects across component subsystems of the PPA (Supplementary Figure 4). In summary, the effect of task modulation on criticality was in general prominent at a finer granularity within functionally defined regions in the domain of vision, suggesting that changes in criticality due to task-related states may construct a hierarchy of criticality that adapts to varying task demands.

## Conclusions

In this study, we sought to establish a connection between brain criticality and cognition. We employed a 2D-Ising model that incorporates anatomical geometry from sMRI data to fit empirical fMRI data obtained during both rest and task. Our findings revealed that while the brain operated at near-criticality, cortical subsystems exhibited heterogeneous changes in criticality depending on cognitive demands, showing a strong coupling between fine-tuning towards criticality and functional engagement. This relationship persisted in a hierarchical fashion, where within the domain of vision, selective fine-tuning occurred in cortical regions specialized in processing different object categories. In summary, these findings suggest that subtle modulation on criticality during tasks may serve as a general mechanism to integrate the functional division of labor within the brain, enabling various cognitive functions.

One fundamental question in neuroscience is how the brain, with its relatively fixed anatomical skeleton, achieves a wide range of cognitive functions. This functional adaptability has been attributed to the variability of neural activity at the microscopic level, where regional neural population activation patterns adapt to accommodate different cognitive demands [43][44][45], and at the macroscopic level, where functional connectivity (FC) among brain regions reconfigures its topological and geometrical properties across tasks [46][47][48]. In this study, we propose a new perspective at the mesoscopic level, subsystems’ criticality, which aligns with task engagement in both the naturalistic task of movie-watching and the well-controlled task of object recognition. Importantly, this new perspective may bridge the conventional microscopic and macroscopic profiles to provide a comprehensive understanding of brain cognition. By finetuning the subsystem criticality, the system could alter the probability of the occurrence of certain configuration (i.e. local activation pattern), and influence the correlation function that in turn induces FC variations among brain regions. Moreover, when operating at or near criticality, the system exhibits high non-linearity of neural responses in both spatial and temporal domains due to the presence of long-range correlations. This could lead to more efficient coding of population patterns [49] and stronger FC magnitudes, thus engendering complex neural behaviors essential for various cognitive processes, thereby bolstering the *cognitive critical brain hypothesis*.

While both resting and task-engaged brains operated at near-criticality, task engagement produced a nuanced shift within the criticality zone, without necessitating drastic phase transitions. Importantly, the shift has two characteristics in terms of location and direction. First, the shift occurred locally rather than globally, akin to a ripple in a pond, primarily observed in subsystems recruited by ongoing tasks. This focused fine-tuning could potentially conserve energy and resources for executing specific tasks while maintaining overall brain functions [50]. Second, the direction of the shift is towards the critical point. Aside from ensuring a more delicately-balanced trade-off between precision and robustness of neural behaviors, the brain at the critical state uniquely capable of integrating and processing information across a broader range of spatial and temporal scales. This leads to optimal information transmission and memory retention, which is required to perform complex cognitive tasks. In summary, the localized and slight shift towards criticality likely enhances the brain’s responses to external stimuli, possibly augmenting its capacity for information integration.

In this study, we treated the brain as a simple, rather than a dynamic system, and therefore utilized statistical mechanics models with clearly defined broken symmetry. This approach allowed us to explicitly measure and comprehend cortical system’s criticality. The finding that the brain operates near Ising criticality implies that the universality of ferromagnetic phase transition could be an effective theory [51] [52] to understand the brain at the large-scale neural population level. This perspective complements previous studies that treat the brain as a complex, far-from-equilibrium dynamic system. Our study thus offers an interpretable perspective on brain dynamics both at rest and during tasks, which could mitigate inconsistencies found in complex dynamic theory measurements (see reviews from [2][50]). On the other hand, the complex dynamic theory may provide additional insights into brain criticality as the brain’s critical state may possess both equilibrium (constant properties over time) and nonequilibrium (variable properties over time) properties. Future research is needed to integrate these perspectives for a more comprehensive understanding of brain criticality.

It is also worth noting that the cortical Ising model diverges from the classic Ising model that describes non-open systems without exhibiting broken time-reversal symmetry [53] [54] [55]. That is, the brain’s Hamiltonian may be non-Hermitian, leading cognitive relevance to the brain’s anisotropy where distinct configurations, or activation patterns, correspond to distinct cognitive processes. Therefore, the transformation of entropy (degeneracy) into negentropy (organization) may facilitate the execution of a wide spectrum of complex functions within a limited range of control parameters. Accordingly, in contrast to classical Hamiltonian systems in nature that typically operate far from the critical state, the brain operates near criticality as revealed in this study. Thus, it is possible that criticality constitutes a distinguishing feature that sets living systems apart from nonliving ones.

Several questions remain unanswered. First, while we found a correlational relationship between criticality and cognition, the underlying causative relationship is unclear. One possible mechanism is that the shift in criticality may change the dimensionality of the neural manifold, transforming representations from being faithful to sensory inputs to those related to decisions and actions [56][57][58]. Future studies employing the topology and geometry may shed light on this connection between neural state space and criticality. Second, while the current study focused on each subsystem, it would be interesting to examine how modulation on criticality might integrate these subsystems into a task-related neural network. Future studies may implement hierarchical structural geometry consisting of long-range connections into Ising models to explore potential scale-dependent variations in brain’s criticality. Finally, because of the observed coupling between criticality and cognition, Ising models may have potential applications in studying neurological and psychiatric disorders, with temperature as a sensitive neural marker.

## Materials and Methods

### Empirical neural data

#### Dataset and participants

Empirical data on brain dynamics was obtained from the S1200 release of HCP-YA dataset [25]. This dataset collected sMRI, rs-fMRI and multiple tfMRI from healthy adult. Informed consent was obtained from all participants with the approval of Washington University Institutional Review Board (IRB) by HCP.

The current study included 3T MRI data from 1018 participants (mean age 28.7 *±* 3.7 years, 546 females), with 98 of them also having 7T MRI data (mean age 29.66 *±* 3.44 years, 57 females). All analyses were performed on the minimally pre-processed MRI data from the HCP dataset. Detailed information regarding the MRI acquisition and the preprocessing pipeline can be found in [25][59]. For convenience, key information is briefly summarized here.

#### sMRI

sMRI Leveraging the emission of electromagnetic signals when tissue hydrogen protons being excited, sMRI including T1-weighted and T2-weighted imaging is able to characterize the volume, geometry, integrity and other anatomical structure properties of brain. The acquired sMRI was first preprocessing, then used to reconstruct 2D brain cortical surface using FreeSurfer. The reconstructed 2D cortical surface contains geometrical and topological information that best suit for sheet-like cerebral cortex. Finally, the reconstructed surfaces in native space of each individual were all registered to MNI standard space and averaged across all participants, resulting a group-level 2D cortical surface. To align with the fMRI spatial resolution, the reconstructed surface with 32k mesh (2 mm resolution) was used.

#### fMRI

While sMRI measures the static structure properties of the brain, fMRI examines the dynamic activity of the brain under specific conditions. fMRI detects the blood-oxygenlevel-dependent (BOLD) signal whose changes are coupled to corresponding local neural activity and therefore allows for the measurement of neural dynamics of the brain in vivo. In fMRI, multiple 3D volumetric frames are initially collected at different time points. These 3D volumetric data were then mapped onto the 2D cortical surface produced by sMRI, resulting in fMRI timeseries data on the 2D cortical surface.

During the rs-fMRI scans, participants were instructed to fixated on a central cross and without any other tasks. For the t-fMRI of movie-watching task, participants were watching concatenated short film clips in the scanners. In the tfMRI of object N-back task, participants were presented with a sequence of pictures and were asked to indicate whether the current picture matched the one presented N-steps earlier. The task was designed using a block design paradigm and included pictures from four categories: face, place, tool, or body, within each block.

The fMRI data, including rs-fMRI (∼15 min per run) and t-fMRI of object N-back task (∼5 min per run), were acquired using a 3T MRI scanner with the following parameters: TR = 720 ms, TE = 33.1 ms, voxel size = 2 *mm*^3^; FOV = 208 × 180 mm, 72 slices, multiband factor = 8.

Another rs-fMRI (∼16 min per run) and t-fMRI of moviewatching task (∼15 min per run) were acquired in a 7T MRI scanner with the following parameters: TR = 1000 ms, TE = 22.2 ms, voxel size = 2 *mm*^3^; FOV = 208 × 208 mm, 85 slices, multiband factor = 5. The current study performed all analyses on the first run of fMRI data for each condition.

#### Parcellation

To characterize the properties of the cortical system at the subsystem level, the MMP based on microstructural architecture, function, connectivity, and/or topography was used [26]. The MMP includes 180 homologous parcels for each hemisphere.

### Cortical Ising model

#### Ising model

For a spin lattice system that each spin can be in either of two states (up or down), the Ising model considers the interactions occurring between neighboring spins that are first-ring adjacent. The Hamiltonian or energy of the Ising model is defined as:

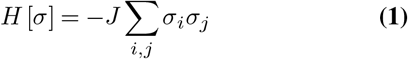

where *J* denotes a constant representing the interaction strength between neighboring spins (set to 1), *σ*_*i*_ denotes the spin state of site *i*, and the summation is over all adjacent spin pairs.

#### Cortical lattice

structure The cortical lattice structure was generated using the reconstructed 2D cortical grayordinates obtained from sMRI. The cortical lattice can be approximated as a triangular lattice, with each vertex (i.e. site) having six neighboring vertices. It is important to note that the cortical lattice was treated as a size-limited lattice without periodic boundary conditions.

#### Monte Carlo simulation

The Wang-Landau algorithm, which is a non-Markovian stochastic process used to estimate the density of system (DOS) states, was applied in this study due to its efficient sampling and fast convergence [28][27]. The simulation was performed on each cortical subsystem (i.e. MMP parcel) at temperatures ranging from 0.001 to 10 with 1000 steps. The cortical lattice was initialized with random spin states (either 1 or -1), and the DOS was set to zero at the beginning. The simulation was run with a flatness criterion of 0.8, an initial modification factor of 1, and a modification factor reducer of 2. The DOS histogram was updated every 105 steps, and the Monte Carlo sweeps continued until the DOS histogram met the flatness criterion, or the modification factor reached 0.0002. Subsequently, the thermodynamic properties of the system, specifically the energy and specific heat, were calculated, resulting in energy and specific heat curves over temperature.

Given that the critical point corresponds to the singularity of the specific heat as a function of temperature, we utilized the peak of the specific heat curve as a feature to identify the critical temperature, and the critical energy was defined as the energy at that temperature.

### Empirical state estimation

#### Fitting empirical neural dynamics to simulation results

The empirical energy and temperature of the cortical Ising model were estimated by fitting empirical neural dynamics provided by fMRI to simulation results. This fitting was performed at the run level for each participant. The fMRI timeseries signal, *s*_*i*_ (*t*), for each vertex *i*, was binarized [60] into up or down states to generate corresponding fMRI spins, *σ*_*i*_ (*t*). This was done using a sign function that utilized the mean of *s*_*i*_ (*t*) across all timepoints as a threshold:

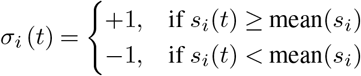

The Ising energy at any given timepoint was calculated from fMRI spins using the Hamiltonian. The median value of the estimated energy across all timepoints in the given run was fitted to the simulation results to obtain the empirical temperature. The Ising energy at any given timepoint was calculated from fMRI spins using the Hamiltonian. The median value of the estimated energy across all timepoints in the given run was fitted to the simulation results to obtain the empirical temperature.

A special within-block binarization was performed for the t-fMRI of the object N-back task to address potential bias in binarization resulting from the block design. The processing was identical to above, except that the binary transformation was carried out in each block with a BOLD signal delay shift of 3 sec.

#### Control analysis with shuffled model

To create a null model for comparison, temporally shuffled fMRI data was generated that only disrupted inter-neuronal interactions while preserving their own statistical information. Specifically, the timeseries of each vertex was independently shuffled across all timepoints within the same run. The resulting temporally shuffled fMRI data was then examined in the same way as the original raw fMRI data.

### Inter-subject correlation

ISC analysis was conducted on the movie watching t-fMRI data to infer the subsystem functional engagement. First, the mean time series was computed for each subsystem across all vertices within it for all participants. To eliminate individual differences in absolute BOLD signal intensity, the data were then standardized by z-scoring along the timepoints. Subsequently, the leave-one-out ISC analysis was conducted by calculating Pearson’s correlation between the fMRI data of each participant and the average of the remaining participants’ data, for each subsystem. The correlation coefficients were then averaged across all participants, yielding a single ISC value for each subsystem.

### Statistical analysis

#### Original data vs. Shuffled data

A paired t-test was conducted to compare the empirical temperature of subsystems between the original and temporally shuffled fMRI data (Fig. 2G). The group-level averaged empirical temperature was used as the dependent variable in the analysis.

#### Correlation between task modulation and ISC

Correlation analysis at the group level was performed utilizing Pearson’s r between task modulation on Ising *Te* (or the original BOLD signal) and ISC across all subsystems (Fig. 3D & E). Task modulation was quantified using t-values from paired t-tests. Specifically, within each subsystem, t-values were obtained by contrasting the Ising *Te* (or average BOLD activation at each vertex) during movie watching with that of the rest condition, for each participant. At the individual level, Pearson’s r was computed between the movie-rest difference ratio and the group-wise ISC across all subsystems (Fig. 3F). The movie-rest difference ratio was defined as (*S*_*movie*_ – *S*_*rest*_) / *S*_*rest*_, where *S* denotes either Ising *Te* or the original BOLD signal.

#### Selective finetuning during object recognition

To examine whether there is heterogenous task modulation in different ROIs, a two-way ANOVA was performed for each hemisphere with the independent variables being ROI and category condition, and dependent variable being the temperature change (Fig. 4C). Specifically, for each participant and each subsystem, the *Te* of each category condition was calculated as the averaged values across all timepoints within the blocks of the given category. The temperature difference of a certain category was then calculated by subtracting category mean *Te* from it. Finally, the temperature change was defined as the ratio of the temperature difference to the category mean temperature.

As a significant interaction effect was found in above ANOVA, simple effect analysis was further conducted to investigate the nature of the effect. One-way repeated ANOVA was performed for each ROI with the factor of category. The statistical results were corrected for multiple comparison with a Bonferroni approach.

## ACKNOWLEDGEMENTS

We would like to express their sincere gratitude to Wenan Guo from Beijing Normal University and Yang Tian from Tsinghua University for providing insightful and valuable comments on the manuscript.

This study was funded by Natural Science Foundation of China (31861143039), Beijing Municipal Science & Technology Commission, Administrative Commission of Zhongguancun Science Park (Z221100002722012), Tsinghua University Guoqiang Institute (2020GQG1016), and Beijing Academy of Artificial Intelligence (BAAI).

## Supplementary Materials

**Supplementary Figure 1.**
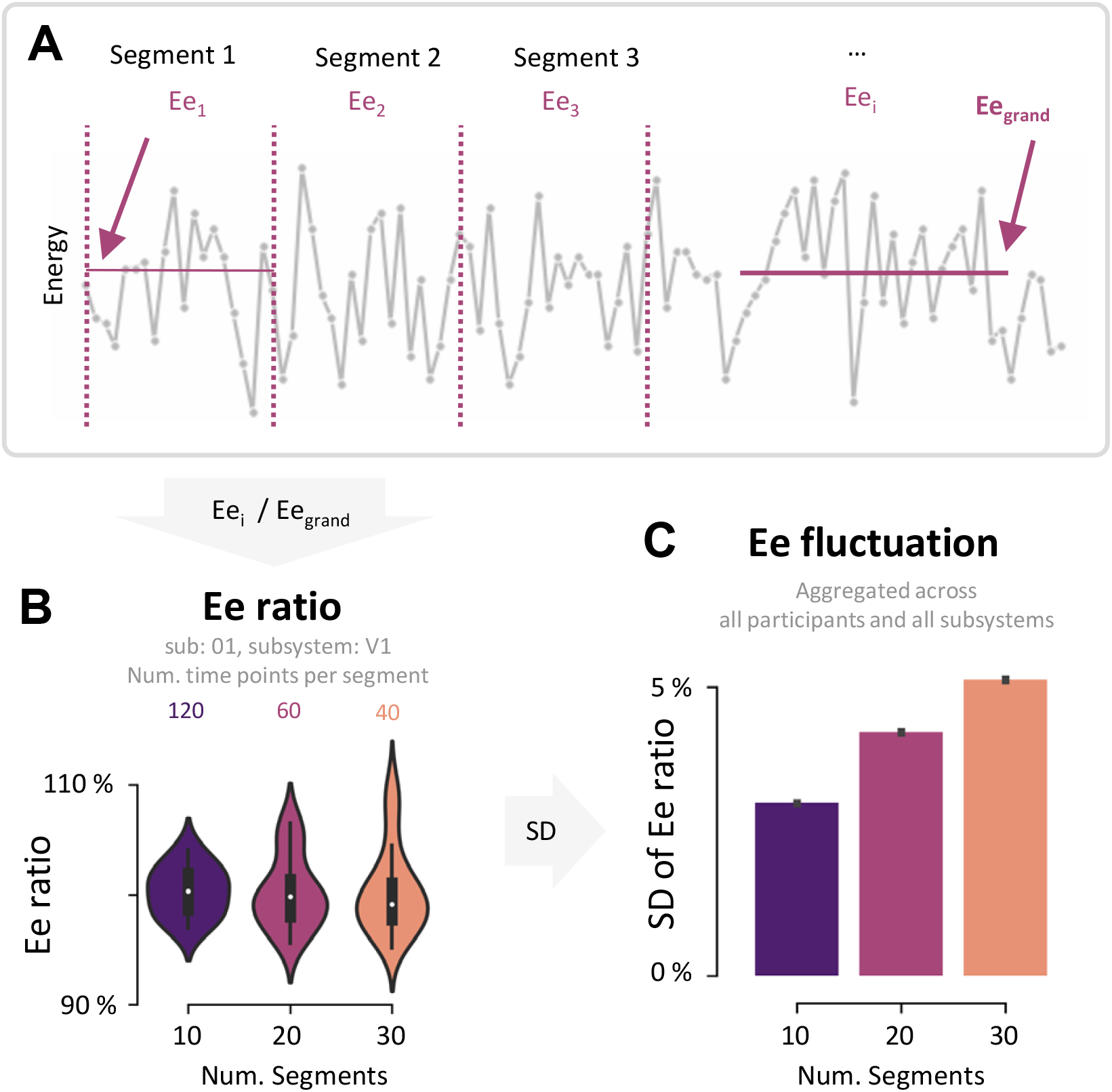
Energy Fluctuation Analysis on cortical subsystems supporting the Ising Model’s Equilibrium Assumption. To test the Ising model’s assumption of system equilibrium, we performed an energy fluctuation analysis on each cortical subsystem. As shown in **(A)**, we divided a run uniformly into 10/20/30 segments and calculated the *E*_*e*_ for each segment *i* (*E*_*ei*_), standardizing it to the grand mean value *E*_*e*grand_ to derive the *E*_*e*_ ratio **(B)**. We then evaluated the variability of the *E*_*e*_ ratio by calculating the SD of *E*_*e*_ ratio values among these segments **(C)**. If the system is in equilibrium, *E*_*e*_ will exhibit minimal variation among different segments. Our data showed all cortical subsystems maintaining slight energy fluctuations (*E*_*e*_ fluctuation *<* 6%) during the entire run, indicating robust system stability and corroborating the equilibrium postulate.

**Supplementary Figure 2.**
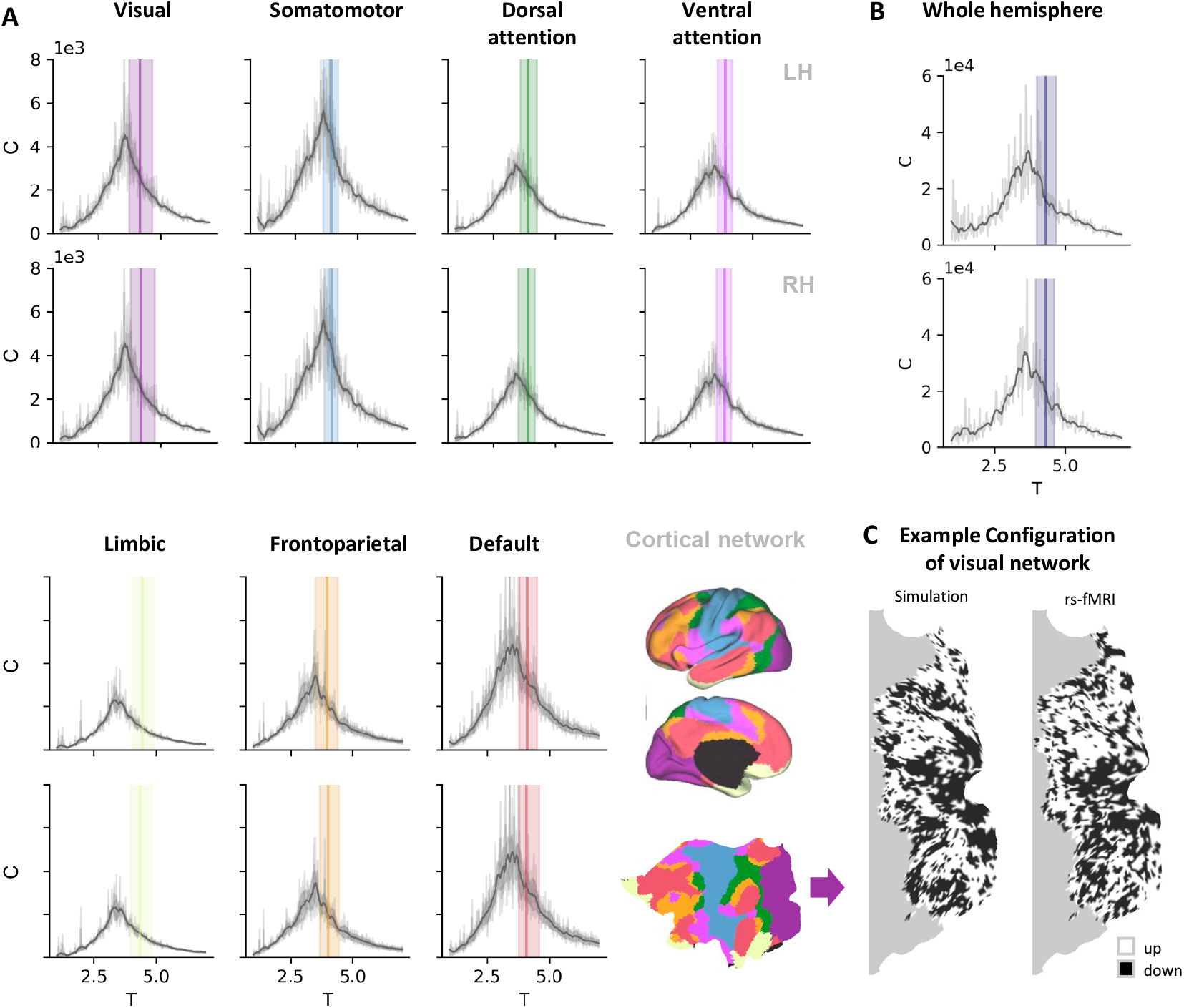
Cortical systems operated near Ising criticality during rest at network and whole hemisphere levels. Similar near-criticality states were also observed at the cortical network [61] level **(A)** and the hemisphere level **(B)**, confirming the near-critical nature of the cortical system across different scales. The conventional Metropolis Monte Carlo algorithm was utilized for numerical simulations in these analyses. Specifically, the temperature range was between 1 and 7, with an interval of 0.02. Equilibration was performed over 2 *×* 10^6^ and 1 *×* 10^7^ steps, and measurements were taken over 5 *×* 10^5^ and 2 *×* 10^6^ steps for networks and whole hemispheres respectively. **(C)** An example configuration of the visual network from the empirical rs-fMRI data and the simulation at the corresponding temperature.

**Supplementary Figure 3.**
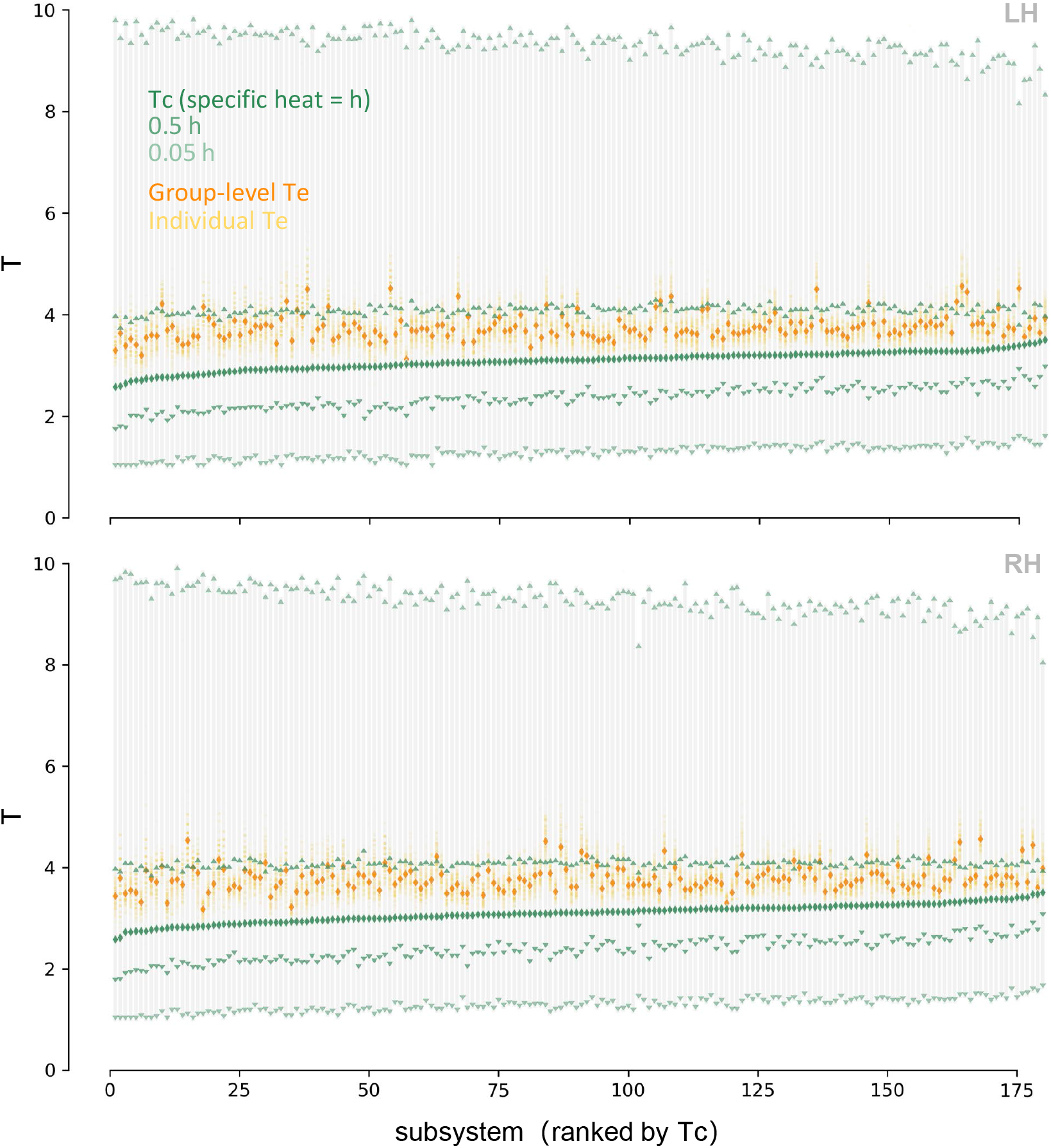
All cortical subsystems operated near Ising criticality during rest (7T rs-fMRI dataset)

**Supplementary Figure 4.**
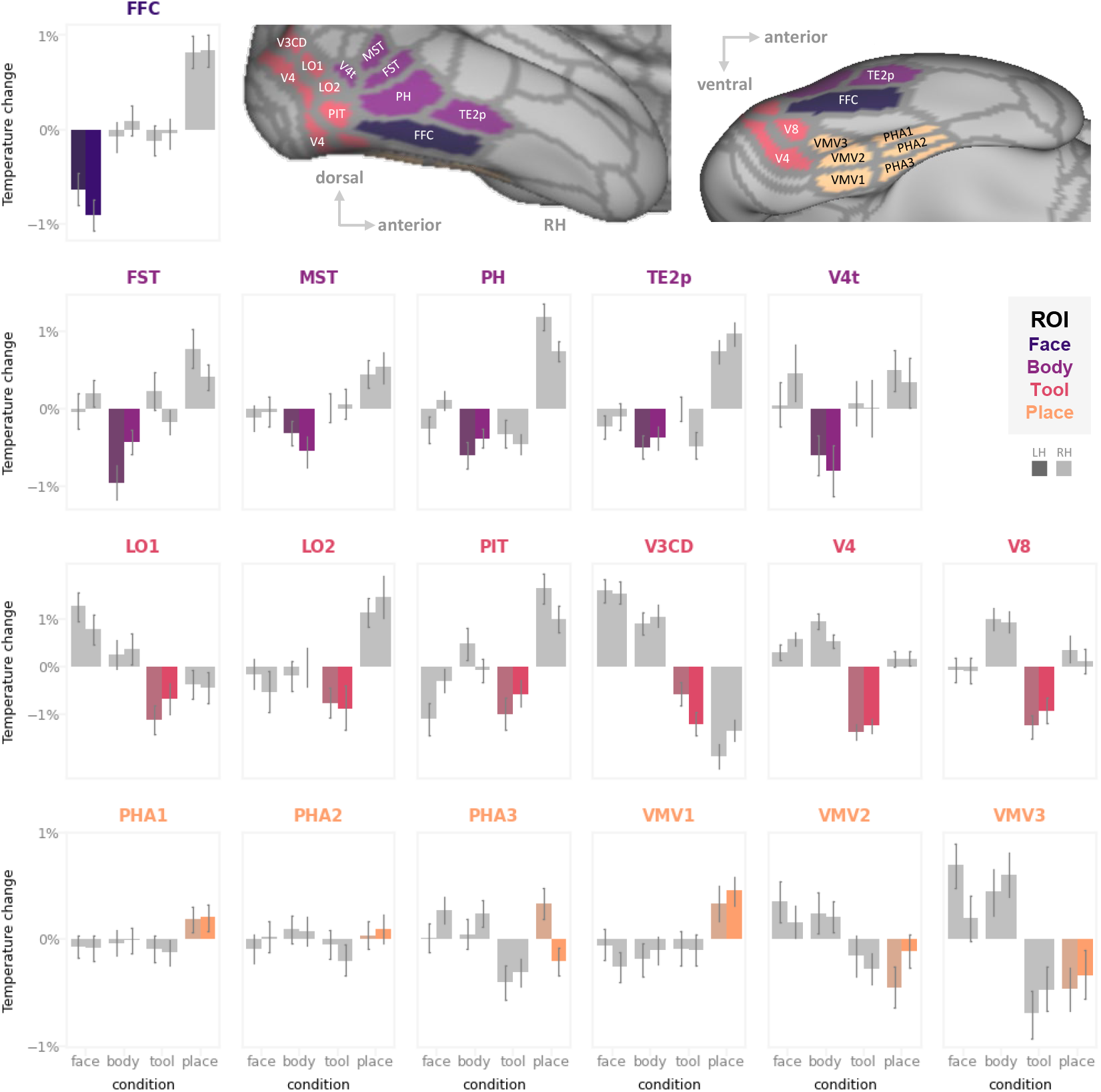
Temperature changes of individual subsystems in the VTC ROIs under different object category conditions.

**Supplementary Table 1.** The cortical subsystems with the highest/lowest movie-rest temperature difference ratio.

| below -2.4% |  |  |  |  |
| --- | --- | --- | --- | --- |
| key | hemi | $(T_{\text{movie}} - T_{\text{rf}}) / T_{\text{rf}}$ | Area Name | Area Description |
| 175 | RH | -3.21% | A4 | Auditory 4 Complex |
| 156 |  | -3.18% | V4t | Area V4t |
| 2 |  | -3.14% | MST | Medial Superior Temporal Area |
| 124 |  | -2.96% | PBelt | ParaBelt Complex |
| 23 |  | -2.89% | MT | Middle Temporal Area |
| 154 |  | -2.85% | VMV3 | VentroMedial Visual Area 3 |
| 107 |  | -2.77% | TA2 | Area TA2 |
| 165 |  | -2.70% | s32 | Area s32 |
| 157 |  | -2.63% | FST | Area FST |
| 173 |  | -2.48% | MBelt | Medial Belt Complex |
| 140 |  | -2.44% | TPOJ2 | Area TemporoParietoOccipital Junction 2 |
| 175 | LH | -3.17% | A4 | Auditory 4 Complex |
| 125 |  | -3.09% | A5 | Auditory 5 Complex |
| 156 |  | -2.85% | V4t | Area V4t |
| 2 |  | -2.85% | MST | Medial Superior Temporal Area |
| 107 |  | -2.78% | TA2 | Area TA2 |
| 173 |  | -2.63% | MBelt | Medial Belt Complex |
| 123 |  | -2.58% | STGa | Area STGa |
| 18 |  | -2.44% | FFC | Fusiform Face Complex |

| above 1% |  |  |  |  |
| --- | --- | --- | --- | --- |
| key | hemi | $(T_{\text{movie}} - T_{\text{rf}}) / T_{\text{rf}}$ | Area Name | Area Description |
| 71 | RH | 1.89% | 9p | Area 9 Posterior |
| 87 |  | 1.84% | 9a | Area 9 anterior |
| 70 |  | 1.47% | 8BL | Area 8B Lateral |
| 72 |  | 1.40% | 10d | Area 10d |
| 86 |  | 1.20% | 9-46d | Area 9-46d |
| 69 |  | 1.12% | 9m | Area 9 Middle |
| 84 |  | 1.04% | 46 | Area 46 |
| 90 |  | 1.01% | 10pp | Polar 10p |
| 87 | LH | 1.56% | 9a | Area 9 anterior |
| 98 |  | 1.53% | s6-8 | Superior 6-8 Transitional Area |
| 70 |  | 1.49% | 8BL | Area 8B Lateral |
| 71 |  | 1.44% | 9p | Area 9 Posterior |
| 72 |  | 1.18% | 10d | Area 10d |
| 68 |  | 1.10% | 8Ad | Area 8Ad |

## References

1. Beggs, J. M. (2008). The criticality hypothesis: how local cortical networks might optimize information processing. Philosophical Transactions of the Royal Society A: Mathematical, Physical and Engineering Sciences, 366(1864), 329–343. URL https://royalsocietypublishing.org/doi/10.1098/rsta.2007.2092

2. Beggs, J. M. (2022). The cortex and the critical point: understanding the power of emergence. Cambridge, Massachusetts: The MIT Press. OCLC: 1333708308.

3. Stein, D. L., & Newman, C. M. (2013). Spin glasses and complexity. Primers in complex systems. Princeton: Princeton University Press.

4. Jensen, O., Kaiser, J., & Lachaux, J.-P. (2007). Human gamma-frequency oscillations associated with attention and memory. Trends in Neurosciences, 30(7), 317–324. URL https://linkinghub.elsevier.com/retrieve/pii/S0166223607001051

5. De las Cuevas, G., & Cubitt, T. S. (2016). Simple universal models capture all classical spin physics. Science, 351(6278), 1180–1183. URL https://www.science.org/doi/10.1126/science.aab3326

6. Beggs, J. M., & Plenz, D. (2003). Neuronal Avalanches in Neocortical Circuits. The Journal of Neuroscience, 23(35), 11167–11177. URL https://www.jneurosci.org/lookup/doi/10.1523/JNEUROSCI.23-35-11167.2003

7. R. Chialvo D. (2004). Critical brain networks. Physica A: Statistical Mechanics and its Applications, 340(4), 756–765. URL https://linkinghub.elsevier.com/retrieve/pii/S0378437104005734

8. Deco, G., & Jirsa, V. K. (2012). Ongoing cortical activity at rest: criticality, multistability, and ghost attractors. Journal of Neuroscience, 32(10), 3366–3375.

9. O’Byrne, J., & Jerbi, K. (2022). How critical is brain criticality? Trends in Neurosciences, (p. 18).

10. Cocchi, L., Gollo, L. L., Zalesky, A., & Breakspear, M. (2017). Criticality in the brain: A synthesis of neurobiology, models and cognition. Progress in Neurobiology, 158, 132–152. URL https://linkinghub.elsevier.com/retrieve/pii/S0301008216301630

11. Yu, S., Ribeiro, T. L., Meisel, C., Chou, S., Mitz, A., Saunders, R., & Plenz, D. (2017). Maintained avalanche dynamics during task-induced changes of neuronal activity in nonhuman primates. eLife, 6, e27119. URL https://elifesciences.org/articles/27119

12. Tomen, N., Rotermund, D., & Ernst, U. (2014). Marginally subcritical dynamics explain enhanced stimulus discriminability under attention. Frontiers in Systems Neuroscience, 8. URL http://journal.frontiersin.org/article/10.3389/fnsys.2014.00151/abstract

13. Shew, W. L., Clawson, W. P., Pobst, J., Karimipanah, Y., Wright, N. C., & Wessel, R. (2015). Adaptation to sensory input tunes visual cortex to criticality. Nature Physics, 11(8), 659–663. URL http://www.nature.com/articles/nphys3370

14. Ma, Z., Liu, H., Komiyama, T., & Wessel, R. (2020). Stability of motor cortex network states during learning-associated neural reorganizations. Journal of Neurophysiology, 124(5), 1327–1342. URL https://journals.physiology.org/doi/10.1152/jn.00061.2020

15. Nonnenmacher, M., Behrens, C., Berens, P., Bethge, M., & Macke, J. H. (2017). Signatures of criticality arise from random subsampling in simple population models. PLOS Computational Biology, 13(10), e1005718. URL https://dx.plos.org/10.1371/journal.pcbi.1005718

16. Priesemann, V., Munk, M. H., & Wibral, M. (2009). Subsampling effects in neuronal avalanche distributions recorded in vivo. BMC Neuroscience, 10(1), 40. URL https://bmcneurosci.biomedcentral.com/articles/10.1186/1471-2202-10-40

17. Clawson, W. P., Wright, N. C., Wessel, R., & Shew, W. L. (2017). Adaptation towards scalefree dynamics improves cortical stimulus discrimination at the cost of reduced detection. PLOS Computational Biology, 13(5), e1005574. URL https://dx.plos.org/10.1371/journal.pcbi.1005574

18. Ponce-Alvarez, A., Jouary, A., Privat, M., Deco, G., & Sumbre, G. (2018). Whole-Brain Neuronal Activity Displays Crackling Noise Dynamics. Neuron, 100(6), 1446–1459.e6. URL https://linkinghub.elsevier.com/retrieve/pii/S089662731830953X

19. Shimono, M., Owaki, T., Amano, K., Kitajo, K., & Takeda, T. (2007). Functional modulation of power-law distribution in visual perception. Physical Review E, 75(5), 051902. URL https://link.aps.org/doi/10.1103/PhysRevE.75.051902

20. Arviv, O., Goldstein, A., & Shriki, O. (2015). Near-Critical Dynamics in Stimulus-Evoked Activity of the Human Brain and Its Relation to Spontaneous Resting-State Activity. The Journal of Neuroscience, 35(41), 13927–13942. URL https://www.jneurosci.org/lookup/doi/10.1523/JNEUROSCI.0477-15.2015

21. Fagerholm, E. D., Lorenz, R., Scott, G., Dinov, M., Hellyer, P. J., Mirzaei, N., Leeson, C., Carmichael, D. W., Sharp, D. J., Shew, W. L., & Leech, R. (2015). Cascades and Cognitive State: Focused Attention Incurs Subcritical Dynamics. The Journal of Neuroscience, 35(11), 4626–4634. URL https://www.jneurosci.org/lookup/doi/10.1523/JNEUROSCI.3694-14.2015

22. Felleman, D. J., & Van Essen, D. C. (1991). Distributed hierarchical processing in the primate cerebral cortex. Cerebral cortex (New York, NY: 1991), 1(1), 1–47.

23. Kanwisher, N. (2010). Functional specificity in the human brain: A window into the functional architecture of the mind. Proceedings of the National Academy of Sciences, 107 (25), 11163– 11170. URL https://pnas.org/doi/full/10.1073/pnas.1005062107

24. Mesulam, M.-M. (1990). Large-scale neurocognitive networks and distributed processing for attention, language, and memory. Annals of Neurology, 28(5), 597–613. URL https://onlinelibrary.wiley.com/doi/10.1002/ana.410280502

25. Van Essen, D. C., Smith, S. M., Barch, D. M., Behrens, T. E., Yacoub, E., Ugurbil, K., Consortium, W.-M. H., et al. (2013). The wu-minn human connectome project: an overview. Neuroimage, 80, 62–79.

26. Glasser, M. F., Coalson, T. S., Robinson, E. C., Hacker, C. D., Harwell, J., Yacoub, E., Ugurbil, K., Andersson, J., Beckmann, C. F., Jenkinson, M., Smith, S. M., & Van Essen, D. C. (2016). A multi-modal parcellation of human cerebral cortex. Nature, 536(7615), 171–178. URL http://www.nature.com/articles/nature18933

27. Wang, F., & Landau, D. P. (2001). Efficient, Multiple-Range Random Walk Algorithm to Calculate the Density of States. Physical Review Letters, 86(10), 2050–2053. URL https://link.aps.org/doi/10.1103/PhysRevLett.86.2050

28. Wang, F., & Landau, D. P. (2001). Determining the density of states for classical statistical models: A random walk algorithm to produce a flat histogram. Physical Review E, 64(5), 056101. URL https://link.aps.org/doi/10.1103/PhysRevE.64.056101

29. Duncan, J. (2010). The multiple-demand (MD) system of the primate brain: mental programs for intelligent behaviour. Trends in Cognitive Sciences, 14(4), 172–179. URL https://linkinghub.elsevier.com/retrieve/pii/S1364661310000057

30. Fedorenko, E., Duncan, J., & Kanwisher, N. (2013). Broad domain generality in focal regions of frontal and parietal cortex. Proceedings of the National Academy of Sciences, 110(41), 16616–16621. URL https://pnas.org/doi/full/10.1073/pnas.1315235110

31. Sonkusare, S., Breakspear, M., & Guo, C. (2019). Naturalistic Stimuli in Neuroscience: Critically Acclaimed. Trends in Cognitive Sciences, 23(8), 699–714. URL https://www.sciencedirect.com/science/article/pii/S1364661319301275

32. Jääskeläinen, I. P., Sams, M., Glerean, E., & Ahveninen, J. (2021). Movies and narratives as naturalistic stimuli in neuroimaging. NeuroImage, 224, 117445. URL https://linkinghub.elsevier.com/retrieve/pii/S1053811920309307

33. Chen, J., Leong, Y. C., Honey, C. J., Yong, C. H., Norman, K. A., & Hasson, U. (2017). Shared memories reveal shared structure in neural activity across individuals. Nature Neuroscience, 20(1), 115–125. URL http://www.nature.com/articles/nn.4450

34. Zacks, J. M., Braver, T. S., Sheridan, M. A., Donaldson, D. I., Snyder, A. Z., Ollinger, J. M., Buckner, R. L., & Raichle, M. E. (2001). Human brain activity time-locked to perceptual event boundaries. Nature Neuroscience, 4(6), 651–655. URL http://www.nature.com/articles/nn0601_651

35. Zadbood, A., Chen, J., Leong, Y., Norman, K., & Hasson, U. (2017). How We Transmit Memories to Other Brains: Constructing Shared Neural Representations Via Communication. Cerebral Cortex, 27 (10), 4988–5000. URL http://academic.oup.com/cercor/article/27/10/4988/4080827/How-We-Transmit-Memories-to-Other-Brains

36. Hasson, U., Nir, Y., Levy, I., Fuhrmann, G., & Malach, R. (2004). Intersubject Synchronization of Cortical Activity During Natural Vision. Science, 303(5664), 1634–1640. URL https://www.science.org/doi/10.1126/science.1089506

37. Nastase, S. A., Gazzola, V., Hasson, U., & Keysers, C. (2019). Measuring shared responses across subjects using intersubject correlation. Social Cognitive and Affective Neuroscience, (p. nsz037). URL https://academic.oup.com/scan/advance-article/doi/10.1093/scan/nsz037/5489905

38. Ohad, T., & Yeshurun, Y. (2023). Neural synchronization as a function of engagement with the narrative. bioRxiv, (pp. 2023–01).

39. Baldassano, C., Chen, J., Zadbood, A., Pillow, J. W., Hasson, U., & Norman, K. A. (2017). Discovering Event Structure in Continuous Narrative Perception and Memory. Neuron, 95(3), 709–721.e5. 00000. URL http://www.cell.com/neuron/abstract/S0896-6273(17)30593-7

40. Cocuzza, C. V., Ruben, S.-R., Ito, T., Mill, R. D., Keane, B. P., & Cole, M. W. (2022). Distributed resting-state network interactions linked to the generation of local visual category selectivity. preprint, Neuroscience. URL http://biorxiv.org/lookup/doi/10.1101/2022.02.19.481103

41. Baldassano, C., Beck, D. M., & Fei-Fei, L. (2013). Differential connectivity within the parahip-pocampal place area. Neuroimage, 75, 228–237.

42. Häusler, C. O., Eickhoff, S. B., & Hanke, M. (2022). Processing of visual and non-visual naturalistic spatial information in the” parahippocampal place area”. Scientific data, 9(1), 147.

43. Haxby, J. V., Gobbini, M. I., Furey, M. L., Ishai, A., Schouten, J. L., & Pietrini, P. (2001). Distributed and Overlapping Representations of Faces and Objects in Ventral Temporal Cortex. Science, 293(5539), 2425–2430. URL https://www.science.org/doi/10.1126/science.1063736

44. Kamitani, Y., & Tong, F. (2005). Decoding the visual and subjective contents of the human brain. Nature Neuroscience, 8(5), 679–685. URL http://www.nature.com/articles/nn1444

45. Norman, K. A., Polyn, S. M., Detre, G. J., & Haxby, J. V. (2006). Beyond mind-reading: multi-voxel pattern analysis of fMRI data. Trends in Cognitive Sciences, 10(9), 424–430. URL https://linkinghub.elsevier.com/retrieve/pii/S1364661306001847

46. Gonzalez-Castillo, J., & Bandettini, P. A. (2018). Task-based dynamic functional connectivity: Recent findings and open questions. Neuroimage, 180, 526–533.

47. Hutchison, R. M., Womelsdorf, T., Allen, E. A., Bandettini, P. A., Calhoun, V. D., Corbetta, M., Della Penna, S., Duyn, J. H., Glover, G. H., Gonzalez-Castillo, J., et al. (2013). Dynamic functional connectivity: promise, issues, and interpretations. Neuroimage, 80, 360–378.

48. Ioannides, A. A. (2007). Dynamic functional connectivity. Current opinion in neurobiology, 17 (2), 161–170.

49. Safavi, S., Chalk, M., Logothetis, N., & Levina, A. (2023). Signatures of criticality in efficient coding networks. bioRxiv, (pp. 2023–02).

50. Girardi-Schappo, M. (2021). Brain criticality beyond avalanches: open problems and how to approach them. Journal of Physics: Complexity, 2(3), 031003. URL https://iopscience.iop.org/article/10.1088/2632-072X/ac2071

51. Weinberg, S. (2021). On the development of effective field theory. The European Physical Journal H, 46, 1–6.

52. Burgess, C. P. (2007). An introduction to effective field theory. Annu. Rev. Nucl. Part. Sci., 57, 329–362.

53. El-Ganainy, R., Makris, K. G., Khajavikhan, M., Musslimani, Z. H., Rotter, S., & Christodoulides, D. N. (2018). Non-hermitian physics and pt symmetry. Nature Physics, 14(1), 11–19.

54. Ashida, Y., Gong, Z., & Ueda, M. (2020). Non-hermitian physics. Advances in Physics, 69(3), 249–435.

55. Gong, Z., Ashida, Y., Kawabata, K., Takasan, K., Higashikawa, S., & Ueda, M. (2018). Topological phases of non-hermitian systems. Physical Review X, 8(3), 031079.

56. Xie, Y., Hu, P., Li, J., Chen, J., Song, W., Wang, X.-J., Yang, T., Dehaene, S., Tang, S., Min, B., et al. (2022). Geometry of sequence working memory in macaque prefrontal cortex. Science, 375(6581), 632–639.

57. Nogueira, R., Rodgers, C. C., Bruno, R. M., & Fusi, S. (2023). The geometry of cortical representations of touch in rodents. Nature Neuroscience, (pp. 1–12).

58. Kaufman, M. T., Benna, M. K., Rigotti, M., Stefanini, F., Fusi, S., & Churchland, A. K. (2022). The implications of categorical and category-free mixed selectivity on representational geometries. Current Opinion in Neurobiology, 77, 102644.

59. Glasser, M. F., Sotiropoulos, S. N., Wilson, J. A., Coalson, T. S., Fischl, B., Andersson, J. L., Xu, J., Jbabdi, S., Webster, M., Polimeni, J. R., Van Essen, D. C., & Jenkinson, M. (2013). The minimal preprocessing pipelines for the Human Connectome Project. NeuroImage, 80, 105–124. URL https://linkinghub.elsevier.com/retrieve/pii/S1053811913005053

60. Watanabe, T., Hirose, S., Wada, H., Imai, Y., Machida, T., Shirouzu, I., Konishi, S., Miyashita, Y., & Masuda, N. (2013). A pairwise maximum entropy model accurately describes resting-state human brain networks. Nature communications, 4(1), 1370.

61. Thomas Yeo, B. T., Krienen, F. M., Sepulcre, J., Sabuncu, M. R., Lashkari, D., Hollinshead, M., Roffman, J. L., Smoller, J. W., Zöllei, L., Polimeni, J. R., Fischl, B., Liu, H., & Buckner, R. L. (2011). The organization of the human cerebral cortex estimated by intrinsic functional connectivity. Journal of Neurophysiology, 106(3), 1125–1165. URL http://www.physiology.org/doi/10.1152/jn.00338.2011

